# Transcription factors mediating regulation of photosynthesis

**DOI:** 10.1101/2023.01.06.522973

**Authors:** Wiebke Halpape, Donat Wulf, Bart Verwaaijen, Anna Sophie Stasche, Sanja Zenker, Janik Sielemann, Sebastian Tschikin, Prisca Viehöver, Manuel Sommer, Andreas P. M. Weber, Carolin Delker, Marion Eisenhut, Andrea Bräutigam

**Affiliations:** Computational Biology, Faculty for Biology, Bielefeld University; Center of Biotechnology, Bielefeld University; Genetics and Genomics of Plants, Faculty for Biology, Bielefeld University; Institute of Plant Biochemistry, Heinrich Heine University Düsseldorf, Cluster of Excellence on Plant Science (CEPLAS), Germany; Institute of Agricultural and Nutritional Sciences at Martin-Luther-University Halle-Wittenberg; Department of Genetics, Martin-Luther-University Halle-Wittenberg

## Abstract

Photosynthesis by which plants convert carbon dioxide to sugars using the energy of light is fundamental to life as it forms the basis of nearly all food chains. Surprisingly, our knowledge about its transcriptional regulation remains incomplete. Effort for its agricultural optimization have mostly focused on post-translational regulatory processes^1–3^ but photosynthesis is regulated at the post-transcriptional^4^ and the transcriptional level^5^. Stacked transcription factor mutations remain photosynthetically active^5,6^ and additional transcription factors have been difficult to identify possibly due to redundancy^6^ or lethality. Using a random forest decision tree-based machine learning approach for gene regulatory network calculation^7^ we determined ranked candidate transcription factors and validated five out of five tested transcription factors as controlling photosynthesis *in vivo*. The detailed analyses of previously published and newly identified transcription factors suggest that photosynthesis is transcriptionally regulated in a partitioned, non-hierarchical, interlooped network.

## Main

Photosynthesis, the process which generates virtually all calories for human consumption, is a very productive and at the same time very dangerous pathway^8^. The photosynthetic electron transfer chain with its photosystems (PSs) and its light harvesting complexes (LHCs) harvests the energy of photons (**Fig. 1A**). This energy needs to be near immediately consumed by the Calvin Benson Bassham cycle (CBBc), photorespiration (**Fig. 1A**), and other reactions to avoid overoxidation and radical production^8^. If the balance between harvest and consumption was only regulated at the post-transcriptional level^4^, large potential savings in protein investment in this high protein investment pathway (**Fig. 1B, SupData1**^9^) were impossible to realize. Therefore, we hypothesized that both pathways are at least partially independently controlled at the transcriptional level.

**Figure 1:**
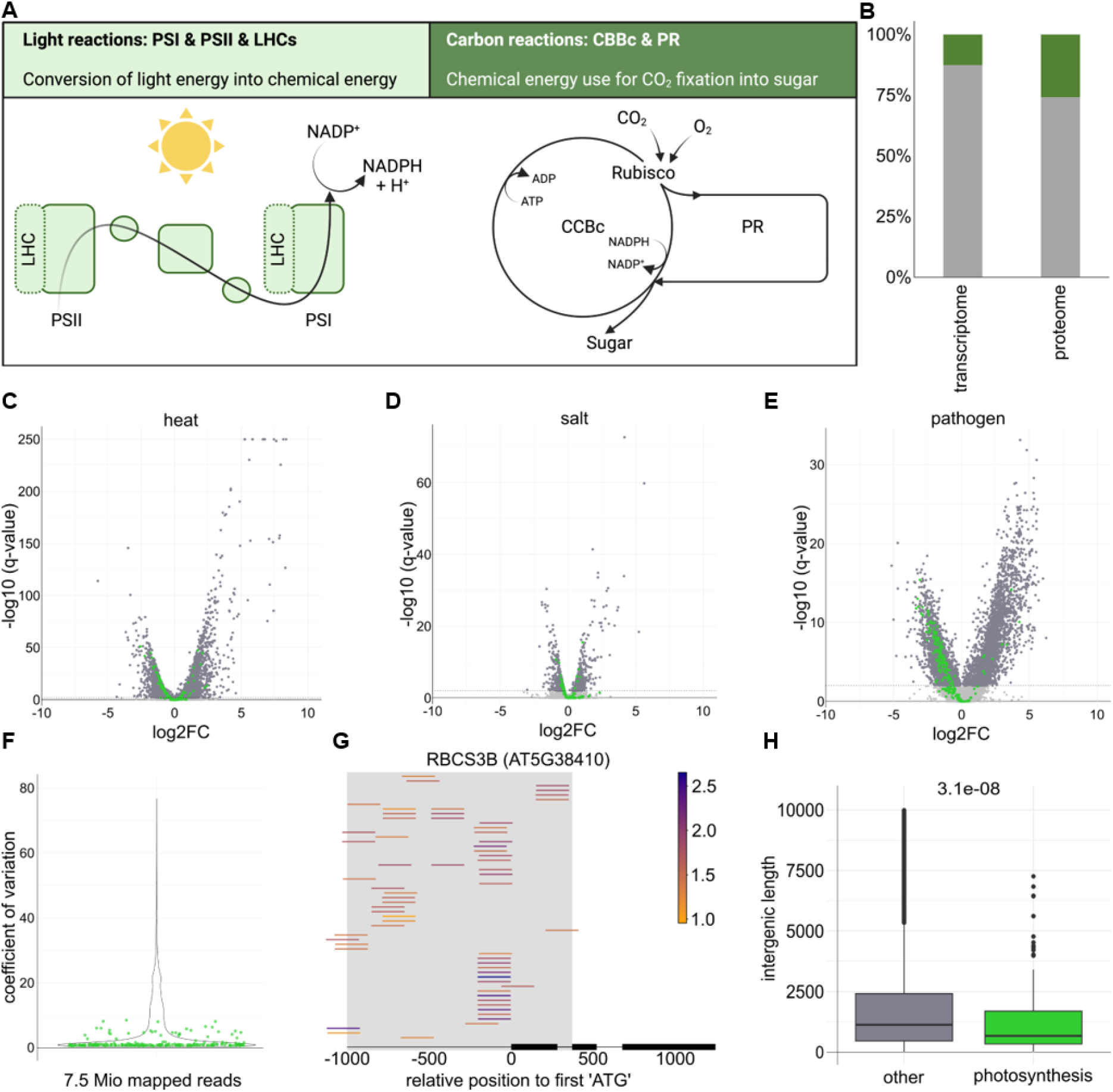
Photosynthetic gene expression is variable. (A) schematic representation of photosynthesis; (B) proportion of *Arabidopsis thaliana* proteome and transcriptome invested in photosynthesis (green); (C-E) volcano plots of transcript abundance changes with photosynthetic genes in green; (F) coefficient of variance for all transcripts in 6,033 RNA-seq experiments, photosynthetic genes in green; (G) experimentally validated binding events on the RBCS3B promoter with relative DAP-seq peak height coded in color; (E) promoter lengths of all promoters vs. photosynthetic promoters (Wilcoxon Rank test, p<1e-7)

The light harvesting module of photosynthesis is at least partially controlled by transcription factors (TFs), which also control germination. Photosynthetic genes are transcriptionally activated by ELONGATED HYPOCOTYL 5 (HY5) ^10^ and repressed by PHYTOCHROME INTERACTING PROTEINS (PIFs) 1, 3, 4, and 5 as evidenced by the *pifq* mutant^11^. Neither mutant was initially identified for their effects on photosynthetic gene expression. Frequently depicted downstream of these regulators^5^, the GOLDEN2-LIKE (GLK) 1 and 2^6,12^ and the GATA-type TFs GATA, NITRATE-INDUCIBLE, CARBON METABOLISM-INVOLVED (GNC) ^13^ and GNC-like^6,14^ of which only the original GOLDEN2 mutation in maize was characterized as a photosynthetic TF^15^. We hypothesized that the identification and characterization of additional photosynthetic regulators will reveal different modes of regulation of photosynthetic modules and will reveal if regulation is hierarchical or more complex.

### Photosynthetic transcript abundance is variable

To understand the regulatory processes related to photosynthesis, we started an in depth analysis of photosynthetic genes and their abundance variation. In mature leaves, photosynthetic transcripts significantly changed in abundance in response to stresses^16^ with changes up to 10-fold (**Fig. 1C to E**) and were among those with the highest absolute changes (**SupFig.1**). Photosynthesis is indeed also controlled at the transcriptional level. To test overall variation, 7,468 RNA-seq experiments in wild type *Arabidopsis thaliana* plants were compiled (**SupData 2**) and 6,033 RNA-seq datasets with at least 7.5 million mapped reads per samples re-analyzed (**SupData 3**). The coefficient of variation for all transcripts was plotted with photosynthetic transcripts highlighted (**Fig. 1F**). Compared to all transcripts, photosynthetic transcripts varied close to the average of all transcripts and were not among those with a large coefficient of variation (**Fig. 1F**). To compile a list of candidate regulators, protein::DNA binding for 217 TFs was analyzed. From 0 to 118 (median = 57) regulators bind to photosynthetic promoters (**SupData 4**, analyzed from^17^). The RbcS promoter, for example (**Fig.1G**), is bound by 57 TFs with many binding to the same sites indicating highly complex regulation, likely with redundant contributions, and a necessity for identifying those TFs with a large contribution to regulation. A detailed analysis demonstrated that photosynthetic promoters were significantly shorter compared to all promoters (p<10^-7^, Wilcoxon Sum Rank Test, **Fig. 1H**) with some as short as 250 bp.

We hypothesized that the variation in photosynthetic transcript abundances (**Fig. 1C-F**) opens the door to RNA-based gene regulatory networks (GRNs) of photosynthetic gene expression. This approach allows to test all TFs for their contribution. Co-expression and linear models have previously been pursued for network construction^18^. We hypothesized that unlike linear approaches, random forest regression decision trees (RF) ^19^ will construct a GRN that includes non-linear and combinatorial interactions between TFs and targets genes and captures direct and indirect interactions. 2,399 TFs were obtained from published sources^20^, curated, and used as regulators for a wrapped GENIE3^19^ algorithm. For validation, biology-dependent evaluation workflows were developed for RF based GRNs (**Fig. 2A**).

**Figure 2:**
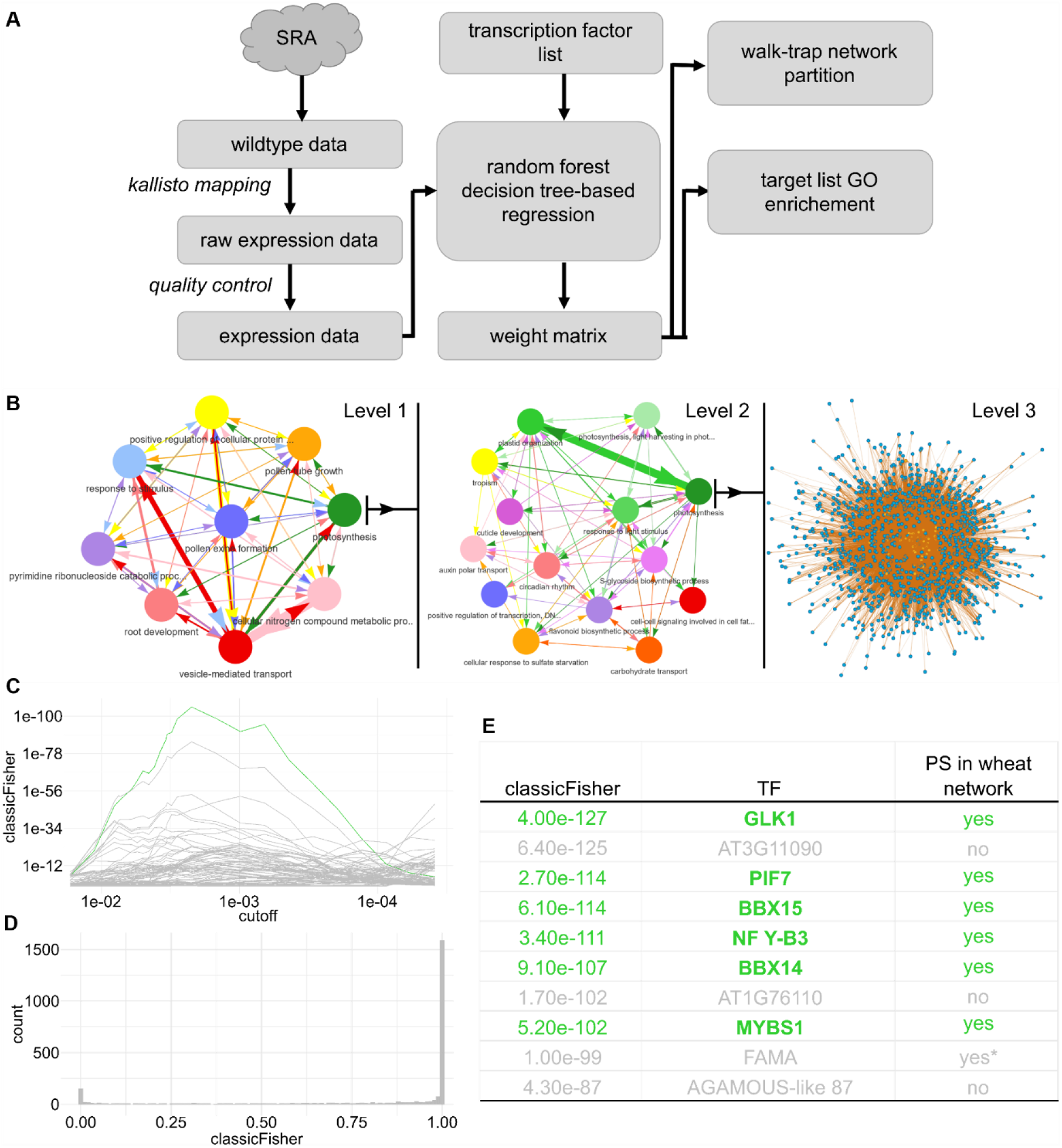
Gene regulatory network based identification of candidate TFs for photosynthesis. (A) scheme of the workflow; (B) iterative walk-trap based partitioning of the RF output and hairball plot of the photosynthetic sub-subcommunity; (C) GO term enrichment for all biological process GO terms for the transcription factor GLK1 at different weight cut-offs in the network; (D) p-value histogram for enrichment of 2,399 TFs in GO:0015979 “photosynthesis”; (E) overlap of ranked list with wheat GRN^21^

### Gene regulatory networks suggest candidates for photosynthetic regulation

The algorithm outputs dimensionless numbers, called weights. These represent the contribution of each TF to abundance values of each target transcript. At a weight cut-off of 0.005, 2,398 of 2,399 TFs were assigned at least one target gene and 33,105 of 33,599 target genes were assigned at least one candidate regulating TF (**Sup.Fig. 2).** The initial publication of GENIE3^19^ and subsequent users^22^ identified a high false positive rate (up to a 80%^19^) at full recall when single edges connecting targets to TFs were considered. This indicates a substantial amount of noise in the data. To test if the communities defined by the GENIE3 output contained biological information, a series of random walks^23^ was conducted on the data (**Fig. 2A, B**). The initial walk-trap detected nine communities of which each enriched for particular gene ontology (GO) terms^24,25^ (**Fig. 2B**). The photosynthesis community was extracted and re-partitioned into 14 subcommunities, which included photosynthesis *per se*, plastid organization, and response to light stimulus enriching subcommunities (**Fig. 2B**). Photosynthesis and plastid organization partitioned into different sub-communities indicating that both processes are at least partially independently regulated, and that the RF based GRN separates both processes (**Fig. 2B**). Plotting the complete network of the photosynthesis subcommunity resulted in 55 candidate TFs (**Fig. 2B**). We tested the community for known photosynthetic regulators and identified GLK1^6^,^12^, GLK2^6^,^12^, GNC^6^,^14^, and GNC-like^6^,^14^ (**SuppData 5**). The GO term enrichments demonstrate that the weight matrix constructed by GENIE3 contained biological information (**Fig. 2B**). The identification of known photosynthetic regulators indicated that regulators of enriched pathways co-localize in communities with their targets. To facilitate analysis of pathways other than photosynthesis, the complete community information is provided as **SuppData 6**. We tested the target genes of each TF for enrichment with a sliding weight cut-off (**Fig. 2A, C**) to obtain a ranked list of photosynthesis regulation candidates to prioritize further validation. We hypothesized that by testing for enrichment of genes in a particular GO term, noise is filtered, and the TF assigned a putative function based on the GO term (**Fig. 2A**). This analysis overcomes the expected high noise level^19^ as the importance of single edges is diminished in favor of significant biological relevance based on many edges. The known photosynthetic transcription factor GLK1^12,26^ was used to test the method (**Fig. 2C**). At different weight cut-offs all GO terms in biological processes were tested for enrichment. At a cut-off of 0.005, the GO:0015979 term “photosynthesis” (**SuppData 7)** enriched with a minimal p-value <1e-100. Other GO terms semantically related to photosynthesis also enriched significantly (**Fig. 2B**). A second peak of enrichment at a lower weight cut-off was observed for many TFs, which contained mostly general GO terms with large membership (**SuppData 8**). All TFs in the dataset were tested for enrichment in photosynthesis at a weight cut-off of 0.005. Enrichment p-values ranged from 7.4E-138 to 1 in a typical significance pattern (**Fig. 2D**) resulting in 95 candidates for regulation of photosynthesis. To select the candidates to enter analysis for nuclear photosynthetic regulation, the list was manually curated from the top. Plastid targeted gene products were excluded from validation analysis as were ZP1 known to function in root hairs^27^, DOT3 known to affect venation patterning^28^, FAMA known to only be expressed in the epidermis^29^, and NCH1 known to interact with phototropins^30^ (**SuppData 10**). The results of a wheat GRN^21^ were overlayed with the remaining top candidates in the Arabidopsis GRN (**Fig.2E**). Six candidates identified as photosynthetic in both wheat and Arabidopsis GRNs were pursued further. The top candidate is again GLK1 (**Fig. 2E**), a known photosynthetic regulator reported to control light harvesting complexes^12,26^. PIF7 is a member of the phytochrome interacting factors^11^, which include the photosynthetic repressors PIF1, PIF3, PIF4 and PIF5^11^. However, PIF7 is not degraded upon light exposure^11^. The B-BOX transcription factors BBX14 and BBX15 are closely related proteins of the BBX family^31^, which also include CONSTANS^31^ and BBX proteins known to interact with HY5^32^. Nuclear factor Y-B3 has been reported as being involved in heat tolerance^33^ and MYBS1 was reported to mediate sugar signaling^34^.

### New photosynthetic transcription factors show partitioned, interlooped regulation

We hypothesized that photosynthetic gene expression is deregulated in plants with changes in expression for their controlling TF. For all candidate photosynthesis regulators, expression data of complementants, knock-out mutants, or overexpression lines were obtained through FAIR data (**Fig. 3A and D**) or RNA-seq data was produced for newly established inducible overexpression lines (**Fig. 3B, C, and E**). Volcano plots showed that photosynthetic genes were significantly deregulated in all lines tested except PIF8 indicating that these TFs indeed control photosynthetic gene expression (**Fig. 3A-E**). Photosynthetic genes were binned to the functions of photosystems and light harvesting complexes (LHCs), Calvin Benson Bassham cycle (CBBc), and photorespiration. Photosynthetic transcript abundance was plotted for new candidates (**Fig. 2E**) and for known photosynthetic regulators to identify shared and different patterns (**Fig. 3F**). Re-analysis of the PIF7 complementant in a *pif4,5,7* knock-out mutant background^35^ showed PIF7 to be a transcriptional activator of the photosystems and LHCs (**Fig. 3F**) in stark contrast to the other PIFs, which strongly downregulate them^11^ (**Fig. 3F**). This observation is in line with PIF7s observed role as an activator^36^. PIF8 showed similar patterns in photosynthetic transcript abundance without any significantly changed photosynthetic genes after 72 hours of induction (**Fig. 3F**). BBX14 decoupled the photosystems and LHCs from photorespiration and the CBBc with repression of the photosystems and LHCs, and induction of the CBBc and photorespiration (**Fig. 3F**) after 72 hours of induction. Permanent induction of either BBX14 or the closely related BBX15 resulted in near white plantlets (**Supp.Fig. 3**) as expected for plants in which the photosystems and LHCs are reduced (**Supp.Data 11**). BBX14 enables the plants to balance light harvesting and processing in the photosystems with consumption of the harvested energy in the CBBc and photorespiration (**Fig. 3F**). Both NF-Y B3 and MYBS1 function as general activators of photosynthesis as indicated by transcriptional effects observed in the constitutive overexpressor of NF-Y B3^33^ and inducible overexpression of MYBS1 (**Fig. 3F**). The effect of permanent NF-Y B3 overexpression was rather small, likely since it requires interactors, such as a NF-Y C and a protein with a CCT domain for its action^37^. Permanent overexpression of NF-Y B3 in wheat supports a photosynthetic role^38^. MYBS1 binds the promoters of most of the significantly increased photosynthetic genes (**Fig. 3G**) pointing toward direct upregulation. The analyses clearly demonstrated that the list of photosynthetic TFs^5^, which currently contains GLK1 and GLK2^6,12,26^, GNC and GNL^6,14^, PIF1,3,4 and 5^11^, and HY5^10^ needs to be amended with PIF7, BBX14 and 15, NF-Y B3 and MYBS1. To see the underlying regulatory network, we also reanalyzed DNA-binding data for HY5, MYBS1^17^, PIF1, PIF3, PIF4, PIF5, PIF5, GLK1, GLK2, GNC and GNL (**Supp. Data 14**). The results demonstrated that the TFs are extensively cross-regulated (**Fig. 3G**). One major integrator of photosynthetic gene expression is the G-box (CACGTG), which is bound by all PIFs and HY5 (**Fig. 3G**). These TFs likely bind with different affinities depending on the particular DNA-shape of the G-box^39^, and with different effects on photosynthetic transcript abundance (**Fig. 3F**). MYBS1-binding is enriched in photosynthetic promoters (Fishers Exact Test, p<1e-21, Fig. 3G) at GATAA^17^. It binds more than 50 photosynthetic genes (**Fig. 3F**) and three photosynthetic regulators (**Fig. 3H**). GLKs bind at GGATT^6,40^ and GNC binds at GATC^6^ and both binding sites are present and occupied in several other TF promoters (**Fig. 3F**) and in photosynthetic promoters (**Fig. 3G, SuppData 12**). Regulation of photosynthetic gene expression is controlled by a complex, non-hierarchical, interlooped network of TFs that frequently bind not only photosynthetic target gene promoters but also each other’s promoters.

**Figure 3:**
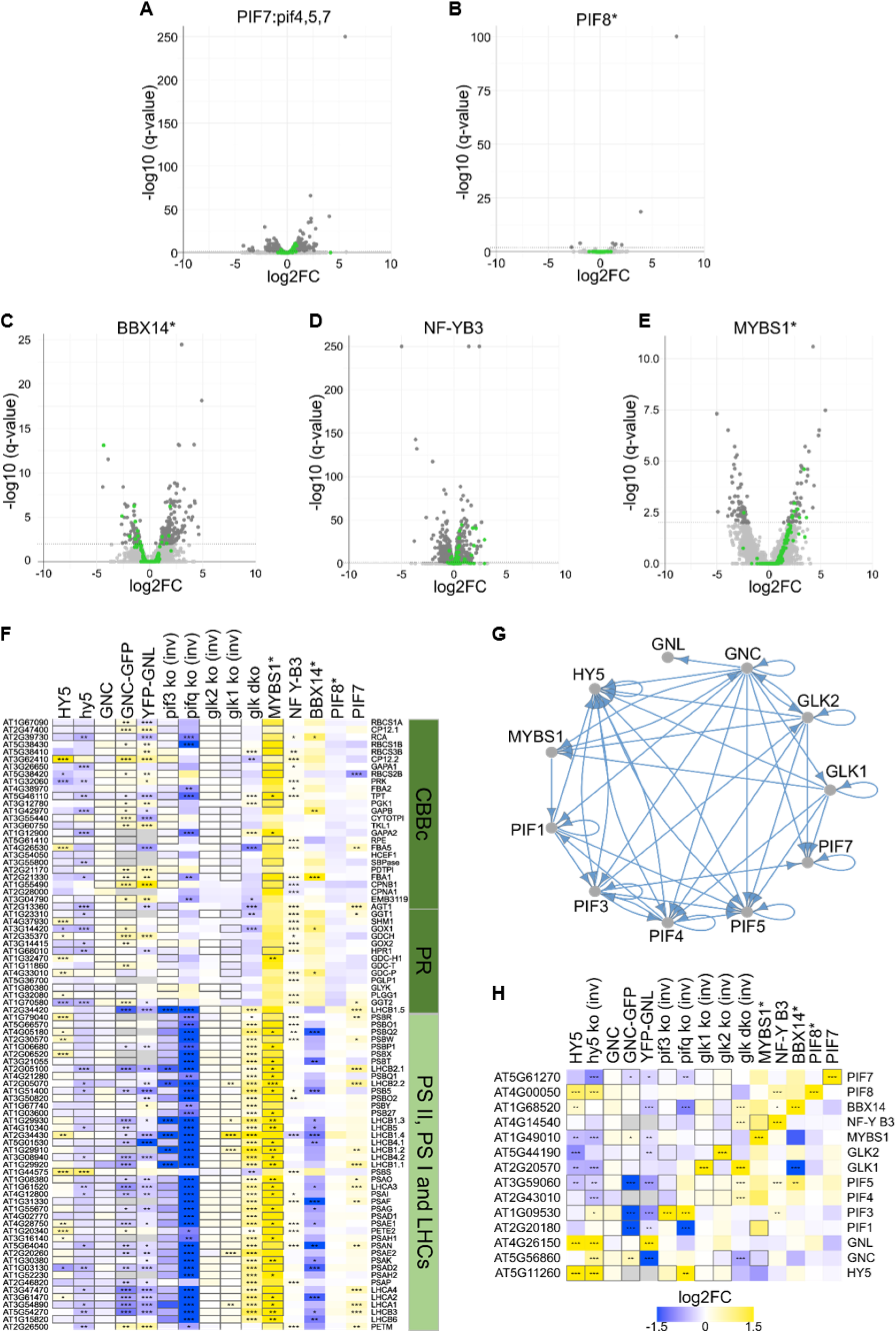
Characterization of photosynthetic TF candidates. (A-E) volcano plots of differential transcript abundance with photosynthetic genes in green for candidate TFs; (F) heat map of transcript abundance changes in photosynthetic genes with significant changes denoted by stars and experimentally validated binding denoted by boxes; starred gene names indicate induced overexpression, mutant data was inverted to homogenize the visualization; (G) diagram of promoter binding data for known photosynthetic TFs; (H) as in F, transcription factors

Despite the highly interconnected regulators, the transcript abundance patterns of photosynthetic genes (**Fig. 3G**) allow classification of the roles for the previously known and newly identified TFs. The *glk1glk2* double mutant identifies GLKs as activators for abundance of photosystem and LHC transcripts (**Fig.3 G, Fig.4**). HY5 also mostly acts as an activator of those although HY5 data is reportedly highly variable between experiments^41^. PIF7 also activates photosystems and LHC genes. In contrast, both NF-Y B3 and MYBS1 activate photosynthetic gene expression throughout while the *pifq* mutant indicates that PIF1, 3, 4, and 5 act as repressors throughout. GNC and GNL partially decouple photosystem and LHC transcript abundance from CBBs and photorespiration transcript abundances while BBX14 fully decouples (**Fig.3 G, Fig.4**). The newly described TFs have sufficient regulatory capacity to up-regulate photosynthetic transcript abundance in general and to decouple the light from the carbon reactions (**Fig. 4**). In concert with the known TFs, general upregulation, general downregulation, upregulation of only light harvesting, and upregulation of only carbon fixation can be accomplished (**Fig. 4**).

**Figure 4:**
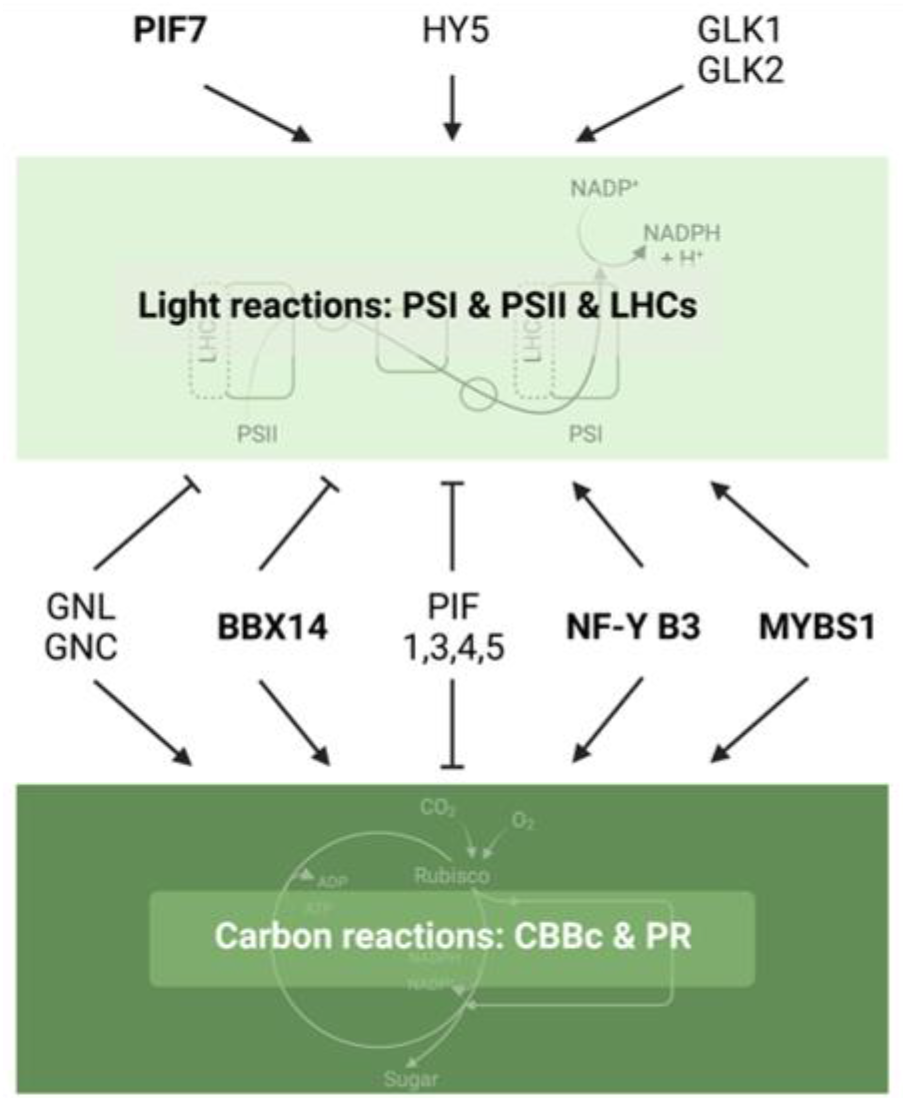
Scheme of photosynthetic gene regulation by a non-hierarchical, interloped network of TFs. The model bases on transcript abundance patterns in mutants and overexpression lines. Dashed lines indicate mixed regulation patterns as depicted in Fig 3G. Newly identified TFs are highlighted in bold.

GRNs calculated using RF, followed by GO term based post-processing, and multi-species comparison are an excellent tool for determining candidate TFs for a specific process (**Fig. 2B and C**, data for all processes available in **SuppTable4, SuppTable5, SuppData2**). The GRNs provided a ranked list of candidate TFs for the process of photosynthesis, all of which except PIF8 could be experimentally validated (**Fig. 3G**). The data demonstrates that photosynthesis is not regulated en bloc, but rather regulated in groups of genes belonging to photosystems and their LHCs (light reaction) and belonging to the CBBc and photorespiration (carbon reactions), respectively (**Fig. 3G, Fig. 4**). The data also demonstrates photosynthesis is regulated through a complex, non-hierarchical, interlooped network of TFs. This more complete list of TFs regulating the different sub-pathways of photosynthesis opens the door to targeted engineering and to a better understanding of the complex outcomes of genetic reconstructions^42^.

## Methods

### Variation analysis

RNA-seq data of ^1^ was downloaded, mapped with kallisto^2^ 0.44 onto the TAIR10 transcriptome and tested for differential expression using edgeR^3^. P-values were Benjamini-Hochberg^4^ corrected and labeled significantly different at p<0.01. 7,048 wildtype RNA-seq datasets were curated from the sequence read archive (SRA) information and downloaded. Each dataset was mapped with kallisto^2^ 0.44.0 in single end (length 200 bp, standard deviation 20) or in paired end mode as specified in SRA. Experiments were filtered for at least 7.5 million mapped reads. The coefficient of variation was plotted for all transcripts as a violin plot with photosynthetic transcripts overlayed in green.

### Promoter analysis

DAP-seq data DNA binding data was downloaded from^5^ and visualized on promoters defined as 1 kb upstream plus sequences including the first intron using a color scale to denote DAP-seq peak height. The intergenic length was determined from the TAIR10^6^ annotation. To test the difference between the intergenic length of the genes with the gene ontology term photosynthesis using a Wilcoxon rank sum test.

### GRN construction and analysis

To infer a gene regulatory network of *A. thaliana* 6033 wildtype RNA-seq datasets (SuppData 13) were downloaded from the sequence read archive (SRA) ^7^ Each dataset was mapped with kallisto^2^ 0.44.0 in single end (length 200 bp, standard deviation 20) or in paired end mode as specified in SRA. Experiments were filtered for at least 7.5 million mapped reads. The tpm values were used as the expression matrix for GENIE3. A manually curated list of DNA-binding transcription factors was derived from TAP-scan^8^. Orthogroups were created using Orthofinder2^9^ using *C. rheinhardtii* v5.5, *V. carteri* v2.0,*M. polymorpha* v3.1, *V. vinifera* v2.1, *S. lycopersicum* iTAG2.4, *A. thaliana* TAIR10, *B. napus* v5, *Z. mays* RefGen v4, *S. bicolor* v3.1.1, *O. sativa* v7.0, *B. distachyon* v3.1, *T. aestivum* IWGSC v1.0, and *H. vulgare* v1 proteins. Proteins with Interpro domains “DNA-binding” and “transcription factor” were added to the curation list. Each entry was manually inspected for functional annotation and those containing the following key words were deselected: cell cycle DNA replication / repair, nucleases, helicases, RNA-binding / splicing, telomere binding, IQ domain and FYVE/PHD-type zinc finger proteins, lipid binding STAR, pentatricopeptide repeats, pleckstrin homolog, protein kinase C-like, protein phosphatase 2C, DUF833. WD40 proteins were kept if they contained CTLH or LisH domains as those were shown to be present in transcription factors. If an orthogroup contained at least one gene annotated as a transcription factor as described above, all orthogroup members were labeled TFs. 2,399 TFs were used as regulators for GENIE3. COP-1 and paralogues included in the list of 2399 transcription factors. For GENIE3 a R C-wrapper was used with the random forest method 1000 trees and the square root of the number of regulators was used for the tree construction. Edges with a feature importance greater than 0.005 were defined as target genes and used for subsequent analysis. For community detection GENIE3 weights were filtered with a 0.005 minimum cutoff and converted to directional edges. Optimized numbers of rounds of iterative community detection were calculated with the walktrap algorithm from the igraph R package (https://igraph.org) with a walk length range 1:20. For each subsequent iteration the walk length resulting in the highest modularity value was picked. Visualizations were created with the visNetwork R package (http://datastorm-open.github.io/visNetwork). The GO-term enrichment for biological process for the target genes of each transcription factor was calculated with the topGO^10^ (V2.5, nodesize=10, classicFisher mode) Rpackage in R 4.2 at different weight cut-offs for the matrix. The GenTable function was modified to report p-values with as small as 1e-1500.

### Single gene analyses

Inducible overexpression lines^11^ were ordered from NASC where available and tested for overexpression in induced plants vs controls using reverse transcription followed by semi-quantitative PCR (data not shown). Overexpression was confirmed using RNA-seq. For BBX14 and PIF8, plantlets were incubated on plates, induced daily with beta-estradiol and harvested after 72 hours of induction. For MYBS1, the coding sequence of MYBS1 with a stop codon was cloned (CGGGCTCAGGCCTGGATGGAGAGTGTGGTGGCAACATG, CCGGGAGCGGTACCCTCAGTGCATTGTCGACGGAGCT) with Gibson assembly between the AscI and XhoI restriction site of the UBQ10:sXVE:HAC:Bar vector^12^. The vector was transformed into *Arabidopsis thaliana* Col-0 using *Agrobacterium tumefaciens* using the floral-dip method^13^. Primary transformed seeds were selected on ½ MS with 25 mg/L glufosinate ammonium. After one week seedlings were picked and placed on soil and were grown in short day conditions for 6 weeks. For each plant two leaves were harvested and vacuum infiltrated 3 times for 20 seconds with 0.2 % ethanol water solution with or without 100 μM β-estradiol. The submerged leaves were put into long day conditions for 12 h. RNA was extracted with the QIAGEN RNeasy Plant Mini Kit with on the column DNAse digest using the QIAGEN RNase-Free DNase. After reverse transcription semi-quantitative PCR with a gene specific and a vector specific primer confirmed the induction of the β-estradiol treated leaves (ATCTGGAACATCGTATGGATACCCGGGAG, ACTGTGAACAATCAAGCTCCTGCGG) and validated via RNA-seq. RNA was isolated using the Qiagen plant RNeasy kit. A sequencing library was constructed with the TruSeq Illumina kit and sequenced on a NextSeq 550. RNA-seq data was further processed as described above.

ChIP and DAP seq data (SuppData 14) was processed using trimmomatic-0.39^14^, aligned to the TAIR10 Arabidopsis genome using bowtie2 version 2.4.1^15^, and filtered for nuclear genome hits, sorted, and converted to bam format with samtools-1.3^16^. For DAP-seq data, peaks were called with GEM version 3.1.4(-d Read_Distribution_default.txt -a 6 -icr 1 -print_bound_seqs -k_min 6 -k_max 20 -k_seqs 600) ^17^ For ChIP-seq data, peaks were called with MACS version 2.2.7.1 (callpeak -gsize 1.118e8 -nomodel -nolambda -keep -dup auto -q 0.05 -call_summits) ^18^. If replicates were available, overlapping peaks were determined^19^ and only peaks present in all replicates kept. A promoter is called bound, if at least one experiment detects TF::promoter interaction at −750 to 0 relative to the transcriptional start site of the target gene. For MYBS1, binding data was downloaded from^5^, processed as above, and enrichment was tested using Fishers Exact Test

### Motif occurrence in photosynthetic promotors

The sequence 750 bp upstream of the transcriptional start site of photosynthesis genes was searched for motif occurrence for the motif of the GLK (GGATT), MYBS1 (GATAA), GNC (GATC) and the G-box (CACGTG) with FIMO^20^. The number of motif occurrences in the promotor was counted per gene.

## Notes

### Competing Interest Statement

The authors have declared no competing interest.

